# Pairing Data Independent Acquisition and High-Resolution Full Scan for Fast Urinary Tract Infection Diagnosis

**DOI:** 10.64898/2026.03.09.710302

**Authors:** Elloise Coyle, Antoine Lacombe-Rastoll, Florence Roux-Dalvai, Mickaël Leclercq, Pascaline Bories, Ève Bérubé, Clarisse Gotti, Dorte Bekker-Jensen, Nicolai Bache, Sandra Isabel, Arnaud Droit

## Abstract

**Background:** Rapid and accurate identification of urinary tract infection (UTI) pathogens is critical for effective treatment and combating antimicrobial resistance. Conventional culture-based diagnostics are slow, and standard tandem mass spectrometry workflows are resource-intensive.

**Methods:** We present a proof-of-concept workflow that integrates high-resolution data-independent acquisition (DIA) MS/MS on the Thermo Scientific Orbitrap Astral with MS1-only spectra from the Orbitrap Exploris 480. DIA data establish a reference panel of pathogen-specific peptides, which are then identified in MS1 spectra from urine samples. Machine learning models trained on these matched MS1 features were used to classify eight common uropathogens and non-infected controls across synthetic inoculations, pure cultures, and clinical patient samples. Model development employed a one-vs-all Random Forest (Ranger) framework with nested cross-validation for feature selection and hyperparameter tuning, followed by evaluation on an independent held-out external patient cohort.

**Results:** The approach accurately distinguished bacterial species in both controlled inoculated samples and clinical patient samples. Using repeated nested cross-validation, the model achieved a mean Matthews Correlation Coefficient (MCC) of 0.88, indicating robust classification performance across resampled training partitions. Performance generalized to an independent patient cohort, achieving an MCC of 0.822, confirming the model’s ability to maintain predictive accuracy under external validation.

**Conclusions:** This proof-of-concept demonstrates that pairing DIA-derived peptide panels with MS1-only data acquired on a cost-effective instrument suitable for routine analysis, enables rapid, culture-free identification of UTI pathogens. The method provides a scalable, high-throughput platform suitable for clinical applications and establishes a foundation for broader biomarker discovery and potential quantitative workflows.

**Key Points:**

- DIA-derived peptide panels enable pathogen detection using MS1-only measurements.
- Machine learning classifies eight common uropathogens covering 84% of infections.
- MS1-only workflows enable culture-free pathogen identification of 300 samples/day.

## Introduction

Urinary tract infections (UTIs) are among the most common bacterial infections globally, affecting millions of individuals each year. It is estimated UTIs affected over 404.6 million individuals in 2019 alone(Zeng *et al*. 2022). From 1990 to 2021, global urinary tract infection cases rose by 66% (from 2.7 to 4.5 billion), deaths nearly tripled to over 300,000, and the total years of healthy life lost due to illness or death (disability-adjusted life years) reached nearly 7 million, underscoring the growing public health burden represented by these infections(He *et al*. 2025). These infections also represent a major economic burden estimated at over US $3.5 billion in the U.S.(Flores-Mireles *et al*. 2015) and €1.5 billion in Europe each year. These infections particularly affect women with about 60 % of whom experience at least one UTI in their lifetime(Guay 2008).

Rapid and accurate diagnosis with precise identification of the causative bacterial species or pathogen is essential for effective treatment of UTIs. This need for effective antibiotic use is especially pressing in the context of rising antimicrobial resistance (AMR). In 2021, over 4.7 million deaths were estimated to be associated with bacterial AMR and projections suggest a continual rise of deaths if AMR continues unchecked(Naghavi *et al*. 2024). Traditionally, UTI diagnosis relies on urine culture, a process that can take 24-72 hours to amplify the bacterial signals to allow further analysis to provide pathogen identification and antibiotic susceptibility. During this time, empirical therapy is often initiated (Brubaker *et al*. 2023). This delay of pathogen identification contributes significantly to inappropriate antibiotic use with a recent study showing 33.4% of UTI prescriptions were suboptimal or inappropriate(Shafrin *et al*. 2022). These inappropriate treatments increase pathogen exposure to antibiotics and increase the chances for bacteria to develop resistance. In this context, there is a critical need to reduce this delay and develop rapid, culture-free diagnostic approaches that can provide same-day results, enabling healthcare professionals to provide treatment tailored to the patient’s needs and reducing inappropriate antibiotic use. Culture-free diagnosis methods are essential to help limit the emergence of resistant strains and improve patient outcomes.

Liquid Chromatography – tandem Mass Spectrometry (LC-MSMS) based proteomics is becoming an increasingly practical clinical tool due to its ability to detect many peptides in complex samples revealing a detailed composition of the different proteins and their origins. It has been successfully used to identify pathogens in urine within a few hours without the need for microbial culture(Roux-Dalvai *et al*. 2019, Gotti *et al*. 2024). By examining protein and peptide patterns, it’s possible to distinguish pathogens, including closely related species of bacteria, and identify markers linked to infection or antimicrobial resistance(Karlsson *et al*. 2020, Deforet *et al*. 2024). In most cases, biomarker discovery through proteomics involves several steps: initial identification using untargeted LC-MSMS, followed by validation and quantitative tracking using targeted methods(Kim *et al*. 2013, Parker and Borchers 2014, Zhu *et al*. 2020, Nakayasu *et al*. 2021). Tandem mass spectrometry is a technique where a sample is analyzed providing an overview of the peptides present in a first spectrum referred to as an MS1 spectrum. Then, either all peptides of the MS1 within specific mass-to-charge ratio (m/z) bins, in the case of Data independent acquisition (DIA), or a certain number of most abundant peptides in the spectra, in the case of Data-dependant Acquisition (DDA), are fragmented and the subsequent fragments are analysed to give a secondary spectrum referred to as an MS2 spectrum. While MS/MS provides detailed information, it can be slow and resource-heavy to be used at scale for numerous biomarkers especially in clinical environments(Grebe and Singh 2011).

More recently, advances in mass spectrometry, notably data-independent acquisition (DIA) methods, have improved both the speed and depth of peptide identification. Instruments like the Orbitrap Astral and Astral Zoom (Thermo Fisher Scientific) including an Asymmetric Track Lossless (Astral) analyzer, allow for rapid, high-resolution profiling in short run times, making them well-suited for biomarker discovery. A recent study showed that the Orbitrap Astral is capable of quantifying five times more peptides per unit than the previous generation Orbitrap instrument operating in data-independent acquisition (DIA) mode(Heil *et al*. 2023). However, these newer machines combining three analyzers (quadrupole, Orbitrap and Astral) come with high cost and operational complexity and are generally inaccessible in a clinical context and are typically limited to research settings.

The simplified Q-Orbitrap type instruments (such as the Orbitrap Q-Exactive, Exploris and Excedion series (Thermo Fisher Scientific)) remain relevant and can provide high resolution spectra albeit with longer runs(Bekker-Jensen *et al*. 2020). These machines have the advantage of being more accessible whilst having high performance in clinical settings(Kardell *et al*. 2023). These machines can perform MS/MS experiments but are limited in the number of peptides that can be detected and fragmented with high resolution in a short run-time when compared to the Astral due to the transient length of the Orbitrap analyzer(Bubis *et al*. 2025). Clinical laboratories for UTI diagnosis must analyse hundreds of samples a day and therefore run-times must be short enough to accommodate the high throughput demands. Whilst fragmentation allows for precise identification of many peptides, short run times decrease accuracy and the 5-minute-or-less run-times required to process 300 samples are too short to acquire MS2 spectra for all of the target peptides required for accurate pathogen identification in urine samples. However, these mass spectrometers are capable of performing high-resolution MS1-only acquisition, which forgoes the fragmentation step and allows for faster, simpler workflows (Liu *et al*. 2007). Nevertheless, without fragmentation and the data provided in the MS2 spectra, peptides can be misidentified or missed in the noisy MS1 spectra which contains many peptide signals(Masselon *et al*. 2008).

Methods exist to exploit the data in MS1 spectra with older techniques such as AMT tagging and more recent predictive methods that attempt to entirely predict the peptide present in the spectra such as MS1direct. The Accurate Mass and Time (AMT) tag method(Pasa-Tolić *et al*. 2004), was introduced in the early 2000s. In AMT, peptides are first identified in a separate experiment using tandem MS and their monoisotopic mass and normalized retention time are stored in a reference library. Experimental samples are then analyzed in MS1-only mode, and observed features (mass and retention time pairs) are matched to the reference library to infer peptide identities. These experiments typically used older methods and did not achieve the depth of modern mass spectrometers.

Recent MS1-only methods such as DirectMS1(Ivanov *et al*. 2020), aim to perform peptide identification directly from MS1 data. DirectMS1 uses predicted peptide properties (mass, retention time, isotope pattern) and fast search algorithms to match experimental MS1 features to a theoretical digest of a protein database. However, this method has a focus on global deep coverage in short runs rather than targeted coverage of specific low-abundance peptides which is required in the case of UTI diagnosis.

In this study, we propose a new method that exploits the capabilities and strength of both the new Orbitrap Astral and the reliable Orbitrap Exploris 480 mass spectrometers by pairing their data and using the detailed in-depth analysis of the Astral’s MS2 spectra to extract precise peptide information from the Exploris’s MS1 spectra. In this proof-of-concept study, we apply this pairing method to provide a mass spectrometry–based workflow for ultra-fast and high-throughput pathogen identification in UTIs that bypasses the need for bacterial culture and the fragmentation step of tandem mass spectrometry analysis. We propose that by using deep peptide profiling via DIA on the Orbitrap Astral to establish a reference panel, we can accurately identify pathogen-specific peptides directly from MS1-only data acquired on the Orbitrap Exploris 480. Here, we focus on applying this method in the context of urinary tract infections (UTIs), focusing on training a machine learning model capable of diagnosing eight of the most prevalent uropathogens in patient data. However, we believe the underlying strategy of matching high-confidence peptide features from MS2 data in high resolution MS1 spectra is broadly applicable to many other contexts necessitating rapid biomarker detection and could be applicable in diverse clinical and diagnostic settings.

### Experimental Procedures

#### Culture of pathogens for inoculations

Microbial strains were obtained from the Culture Collection of Centre de Recherche en Infectiologie of Université Laval (CCRI) registered as WDCM861 at the World Data Centre for Microorganisms. The microbial strains used and their corresponding culture conditions are listed in (**Supp.Table 1**). Each bacterial strain was plated on blood agar (TSA II 5% Sheep blood; Becton Dickinson) and yeast strains were plated on Sabouraud Dextrose Agar (Becton Dickinson) incubated at 35 °C overnight under aerobic conditions (or in air + 5% CO_2_ for *S. agalactiae*). Microbial colonies were then collected and resuspended in phosphate-buffered saline (PBS, 137 mM NaCl/6.4 mM Na_2_HPO_4_/2.7 mM KCl/0.88 mM KH_2_PO_4_ at pH 7.4) at an optical density of 0.5 McFarland.

Thirty microliters of this suspension (hundred microliters for *C. glabrata.)* were used to inoculate 3 mL of BHI medium (Brain Heart Infusion medium broth; Becton Dickinson) or 3 mL of YPD medium (Yeast peptone Dextrose broth) which was incubated overnight at 35 °C under aerobic conditions with agitation (or 35 °C in air + 5% CO_2_ without agitation for *S. agalactiae*). Finally, broth cultures were done by inoculation of 30 μl (100 μL for *C. glabrata)* of the previous culture in 3 mL BHI or YPD medium and incubated in the same conditions. The cultures were stopped in an exponential phase according to the previously known semi-log growth curve of each strain (**Supp.Table 1**). Microbial cultures were counted by incubation of 50 uL of serial dilutions on Blood agar plates.

#### Urine collection

Urine specimens from patients (reported negative or positive by the gold standard method including microbial culture followed by MALDI-TOF analysis) were collected at the CHU de Québec - Université Laval hospital, according to an ethical protocol approved by the Comité d’Éthique de la Recherche du CHU de Québec - Université Laval (recording number 2016-2656). Only midstream urines with a minimum volume of 20 mL were included. Collected urines were aliquoted in 1 mL volume (Eppendorf tube). All samples of urine were stored at -80°C.

#### Protein extraction and enzymatic digestion

For inoculated samples, each pathogen inoculation was prepared by addition of a volume of culture equivalent to 10^6^ CFU based on microbial count in one milliliter of negative urine. For clinical samples, 1 mL of urine specimen was used. Then both types of samples were processed and analyzed the same way, with the exception of a rapid centrifugation (1 min at 100 g) that was added at the beginning of the process of clinical samples to remove large human cell debris. Microbial cells were then isolated by centrifugation at 10,000 g for 10 min. The pellet was washed with 900 µl of 50 mM Tris centrifuged for 10 min at 10,000 g and the supernatant was discarded. Cell lysis was performed by resuspension of the pellet in 50 mM ammonium bicarbonate containing 10 units of mutanolysin enzyme (Mutanolysin from Streptomyces globisporus ATCC 21553, Sigma-Aldrich). Samples were transferred to a 96-well-plate and incubated at 37 ◦C for 1 hour (500 rpm). Bacterial cells inactivation and protein denaturation were then achieved by the addition of 1% sodium deoxycholate (SDC) and 20 mM DTT (dithiothreitol) for 10 min and heating at 95◦C. Proteins were then digested using 250 ng of trypsin enzyme (Sequencing Grade Modified Trypsin, Promega) and incubated for 1 h at 37◦C (500 rpm). Samples were acidified with 5 μl of 50% formic acid (FA) to stop digestion and precipitate the SDC. After 15 min of centrifugation at 4,000 g, peptides contained in the supernatant were transferred in 96 well-plates, vacuum-dried and stored at −80◦C until mass spectrometry analysis.

### Experimental Design and Statistical Rationale

Multiple datasets were used to train and evaluate our machine learning model including two batches of inoculated urine samples from donors deemed negative by the gold-standard method and two batches including clinical specimens collected from patients presenting positive for a urinary tract infection by gold-standard testing. The inclusion of multiple datasets allowed our model to be robust when faced with potential variations between donor urines and batches.

#### Urine Inoculation samples

Urine samples were collected across multiple batches to train and evaluate bacterial classification models. A first batch consisted of 216 samples prepared by inoculating negative donor urines with reference strains of eight target uropathogens (*Escherichia coli*, *Enterococcus faecalis*, *Klebsiella pneumoniae*, *Pseudomonas aeruginosa*, *Streptococcus agalactiae*, *Staphylococcus aureus*, *Staphylococcus saprophyticus*, and *Proteus mirabilis*) at a concentration of 10⁶ CFU/mL, alongside 24 uninoculated negative samples. Donor urines and bacterial strains were randomly paired to introduce biological variability and minimize background biases. These samples were used for both biomarker identification and model training.

A second batch included 623 samples spanning 12 pathogen species, with the eight target bacteria and four additional species (*Acinetobacter pittii*, *Candida glabrata*, *Serratia marcescens*, and *Providencia stuartii*) inoculated at high (10⁶ CFU/mL) and low (10⁵ CFU/mL) concentrations. After filtering to retain only samples of the eight target bacteria and negative urine samples (blanks), 47 negative samples, 24 low-concentration samples and 24 high-concentration samples per class remained for training. Samples containing non-target pathogens were held for testing model behavior on unknown species.

#### Bacterial Culture samples

Additionally, 48 of the pure culture samples previously prepared to create the inoculations (six per bacterial species) were included in machine learning training to reinforce species-specific signal features and improve model performance.

#### Clinical Urine Specimens

A first batch of clinical urine samples were collected from 119 patients at the CHUL outpatient clinic between December 2023 and March 2025. Of these, 54 samples contained target species that were included in training, while 15 samples containing non-target or polymicrobial infections were reserved for exploratory testing. The 50 remaining samples were negative (blank) samples. Negative samples and samples containing target pathogens were included in machine learning training to represent urine and strain heterogeneity of clinical samples.

A second batch of clinical urine samples were collected from 127 patients at the CHUL outpatient clinic also between December 2023 and March 2025. Of these, 98 samples contained target species that were held out for external validation, while 29 samples containing non-target or polymicrobial infections were reserved for exploratory testing.

### Mass spectrometry acquisition

#### DIA analyses

Peptide digestion extracts of samples from the first batch were injected on a nanoLC/MSMS (nanoflow Liquid Chromatography - tandem mass spectrometry) system. The experiments were performed with an Evosep One (Evosep Inc., Odense, Denmark) liquid chromatography coupled to an Orbitrap Astral mass spectrometer (Thermo Fisher Scientific, San Jose, CA, USA), in Data-Independent Acquisition (DIA) mode. 500 ng of peptides loaded on Evotip Pure devices (Evosep Inc., Odense, Denmark) according to the manufacturer protocol were separated using the pre-programmed 300 samples per day method (3.2-minute gradient) on an EV1107 capillary column (4 cm length, 150 µm I.D., 1.9 µm beads, Evosep Inc.) at ambient temperature. Spray voltage was set to 1,900 V. DIA mass spectra were acquired in the Astral analyzer with 200 DIA windows of 3 m/z and no overlap covering a precursor range of 380 to 980 m/z. The average cycle time was approximately 370 ms corresponding to 7-8 points per chromatographic peak. Maximum injection time was set to 3.5 ms, HCD collision Energy to 25 % and AGC Target to 500 %. Orbitrap resolution for MS1 scans were set at 180,000. Internal calibration using fluoranthene radical ions provided by EASY-IC source was used.

#### MS1 Full scan analyses

The same samples were also injected on an Evosep One (Evosep Inc., Odense, Denmark) liquid chromatography system interfaced to an Orbitrap Exploris 480 mass spectrometer (Thermo Fisher Scientific, San Jose, CA, USA) operating in Full Scan (FS) MS1 acquisition mode. 500 ng of peptides were loaded on Evotip Pure devices and separated using the 300 SPD method on EV1107 capillary column. MS1 spectra were acquired in the Orbitrap detector with a 180,000 resolution and a scan range from 300 to 1,500 m/z. AGC target was set to 300 % and maximum injection time was in Auto mode. Samples from the second batch of inoculations and clinical urine specimens were analyzed solely on the Exploris 480 following the same Full Scan acquisition method.

For all mass spectrometry experiments, the samples were injected in a randomized order using simple randomization.

### Mass spectrometry analysis

Peptide identification and quantification from DIA raw files was obtained using DIA-NN(Demichev *et al*. 2020) software (version 1.8.1). This software was first used to generate predicted library from Uniprot FASTA file of each bacterial species of interest and for *Homo sapiens* (**Supp.Table 2**), DIA-NN default parameters were used. Then, each predicted library was used to search against the raw files corresponding to inoculation of the corresponding species. Negative sample files were searched with the *Homo sapiens* library. For all searches, Trypsin/P enzyme parameter was selected with one possible missed cleavage. Carbamidomethylation of cysteines was set as fixed modification. Methionine oxidation, acetylation of protein N-terminus and N-terminal Methionine excision were set as variable modifications. DIA-NN default mass search tolerances were used. Maximum precursor False Discovery Rate (FDR) was set at 1%. The ‘match between runs’ option was used and the protein inference was based on protein names, in non-heuristic mode. The quantification strategy was set to Robust LC mode. All the other parameters were set at default values. The DIA-NN outputs of each search were then aggregated at precursor level using R software(Ihaka and Gentleman 1996, R Core Team 2020) and the following filters were applied : all the precursors identified in negative samples were removed and only precursors found in at least 75% of the replicates for each bacterial species were kept in order to retain the best peptides for each bacteria species. Retention time, m/z, and intensity values were extracted to create a reference set of candidate biomarkers. Complementary isotope information (M, M+1, M+2) for all candidate biomarkers was obtained using Skyline(Pino *et al*. 2020).

### R-based preprocessing and feature extraction

Orbitrap Exploris MS1 data were processed in R (Beagle Scouts version 4.3.1)(R Core Team 2020) to generate model-ready matrices **(Supp.Fig 1)**. All mzML files were converted to CSV format using the Spectra library(Rainer *et al*. 2022), with large tables handled using data.table(Barrett *et al*. 2025). Extracted m/z, intensity, and retention time values were matched to the reference set of candidate biomarkers obtained from DIA analysis within ±3 ppm mass tolerance using raw_acquisition_to_targets. Retention time filtering ±5 seconds around the expected RT was applied via get_rtimes. Isotope clusters containing at least M, M+1, and M+2 signals were retained using get_isotope_presence. For each precursor, the maximum intensity within the RT window was summarized using process_max_intensity_summary. The resulting feature-by-sample matrix included sample IDs and class labels for downstream machine learning. Data manipulation and string matching was performed using tidyverse packages(Wickham *et al*. 2019). Two CSV input files were required: one defining expected m/z and RT from DIA runs and one providing isotope m/z values with ±3 ppm windows from Skyline(Pino *et al*. 2020).

### Model input and feature harmonisation

For machine learning, samples of the eight target pathogens from the first and second urine inoculation batches, along with pure bacterial cultures of the same species and one batch of clinical specimens, were used. Non-target pathogens were excluded from model training but retained for exploratory evaluation as edge cases. One entire batch of clinical urine specimens was held out entirely for external validation.

Feature harmonisation was performed by constructing a unified feature matrix across all datasets. All DIA-derived precursor features were retained, and datasets were merged into a single matrix. Missing feature values were set to zero to ensure a consistent feature space across all samples.

### One-vs-all Random Forest classification

Classification was performed using a one-vs-all Random Forest framework implemented with the ranger package in R(Wright and Ziegler 2017). For each of the eight bacterial classes, an independent binary classifier was trained to distinguish the target class from all remaining classes as well as a classifier to distinguish negative samples (blank) **(Figure 1).**

**Figure 1:**
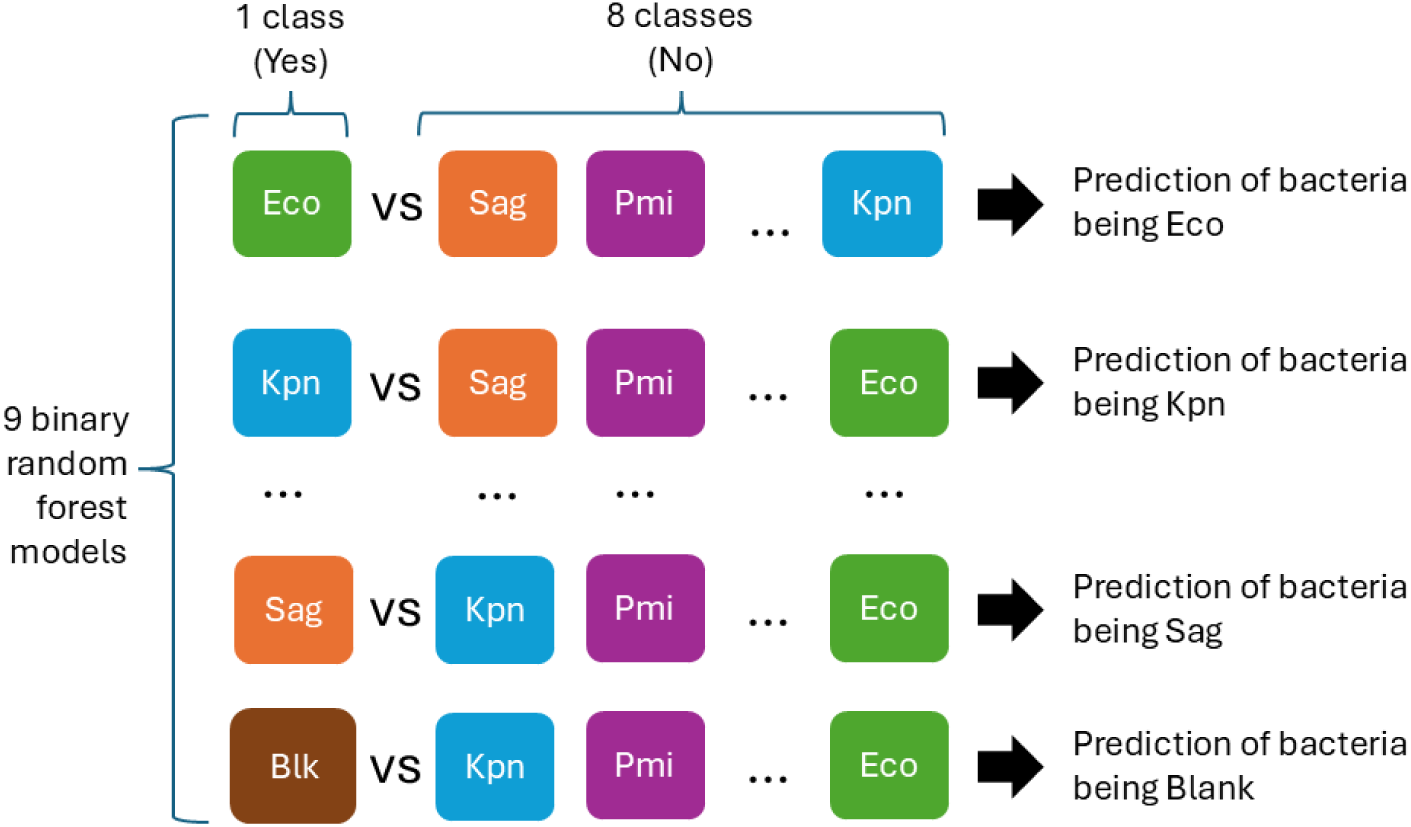
Illustration of One vs All model structure used. Nine binary Random Forest classifiers were trained in a one-vs-all manner to distinguish each target class: Escherichia coli (Eco), Enterococcus faecalis (Efa), Klebsiella pneumoniae (Kpn), Pseudomonas aeruginosa (Pae), Streptococcus agalactiae (Sag), Staphylococcus aureus (Sau), Staphylococcus saprophyticus (Ssa), Proteus mirabilis (Pmi), and blank (Blk, representing non-infected samples) from all other classes. Each model outputs a score representing the probability of a sample belonging to each class.

Binary labels were defined as “Yes” for the target class and “No” for all other classes. For each sample, class probabilities were generated across all one-vs-all models, and final class assignment was performed using a winner-takes-all approach based on the highest predicted probability.

A nested cross-validation framework was used for hyperparameter tuning and feature selection. The outer loop consisted of repeated stratified 70/30 train–test splits across 20 random seeds, with all model development restricted to the outer training set. The outer test set was used only for final performance evaluation.

Within each outer training set, feature selection and hyperparameter tuning were performed using a 5-fold inner cross-validation. Feature selection was conducted separately for each one-vs-all classifier using permutation-based variable importance from Random Forest models (*ranger*). For each class, the top-ranked 100 features were extracted within each inner fold, and a consensus feature set was derived by retaining features that consistently appeared across folds above a predefined frequency threshold (3 folds).

Hyperparameter optimization (mtry fraction, minimum node size, and number of trees) was performed within the same inner cross-validation loop. For each hyperparameter combination, one-vs-all Random Forest models were trained using the consensus feature sets and evaluated on the corresponding inner validation folds using the Matthews correlation coefficient (MCC). The parameter set yielding the highest mean MCC across inner folds was selected.

### Evaluation of model performance

Model performance was quantified using Matthews Correlation Coefficient (MCC) as the primary metric. Per class specificity, sensitivity and balanced accuracy were also calculated using Caret(Kuhn, and Max 2008). Multi-class predictions were derived using the winner-takes-all probability approach. Generalisation and clinical performance were evaluated on the independent external clinical cohort.

### Evaluation of model stability

Model stability was assessed using performance variability across the 20 outer cross-validation iterations. Feature selection stability was quantified by recording feature occurrence frequency across outer folds. The top 50 most frequent features per class across outer iterations were defined as stable and retained for the final one vs all models.

### Evaluation on edge cases

The final models were further assessed on inoculations with unknown pathogens (*Acinetobacter pittii* (Api), *Candida glabrata* (Cgl), *Serratia marcescens* (Sma), and *Providencia stuartii* (Pst)) at high concentrations (10^6^ CFU/mL) and low concentrations (10^5^ CFU/mL), and on previously excluded patient samples containing unknown species or coinfections.

### Model interpretability and confidence

Prediction confidence was evaluated using the maximum class probability for each sample. Classification uncertainty was assessed using the probability margin, defined as the difference between the highest and second-highest predicted class probabilities.

These measures were computed for all outer test folds and external validation samples.

### Software and implementation

All analyses were performed in R. Data preprocessing and feature alignment were implemented using custom scripts.

Machine learning was performed using the ranger package. MCC performance metrics were computed using mltools(Gorman 2021).

A Shiny application was developed to integrate feature extraction, preprocessing, and classification. Analyses were parallelised using furrr(Vaughan, Bengtsson, and Dancho 2026) for computational efficiency, with multisession execution across available cores.

## Results

In this study, we present a proof-of-concept workflow for the rapid, culture-free identification of urinary tract infection pathogens on a Q-Orbitrap instrument suitable for routine analysis. We paired high-resolution DIA data acquired on an Orbitrap Astral with fast MS1-only mass spectrometry data acquired on an Orbitrap Exploris, then trained a machine learning model to classify urinary pathogens from matched MS1 features in five-minute runs. Our experimental workflow relies on sample sets containing different types of samples : i) pure cultures of the eight pathogens most frequently found in UTIs; ii) negative donor urines inoculated at two different levels of concentrations with the same pathogens and four non-target non-UTI related pathogens used as controls; iii) uninoculated negative urines used as controls; iv) clinical samples that were directly collected from patients for final testing (**Figure 2**). A first batch was analyzed in DIA and used to generate a reference set of pathogen-specific peptides, while all subsequent inoculations and clinical specimens were measured using rapid MS1-only acquisition. Model training sets consisted of MS1-only data from inoculations, clinical samples and pure bacterial cultures to classify eight common uropathogens and negative (blank) samples. Model performance was evaluated across outer-iterations of the nested cross-validation framework and validated on an independent clinical cohort. Model behaviour was also observed on non-target pathogens that were retained as “edge cases” (**Figure 2)**. This approach provides a scalable, culture-free workflow for rapid pathogen identification in urine of 8 primary pathogens covering 84% of urinary tract infections.

**Figure 2:**
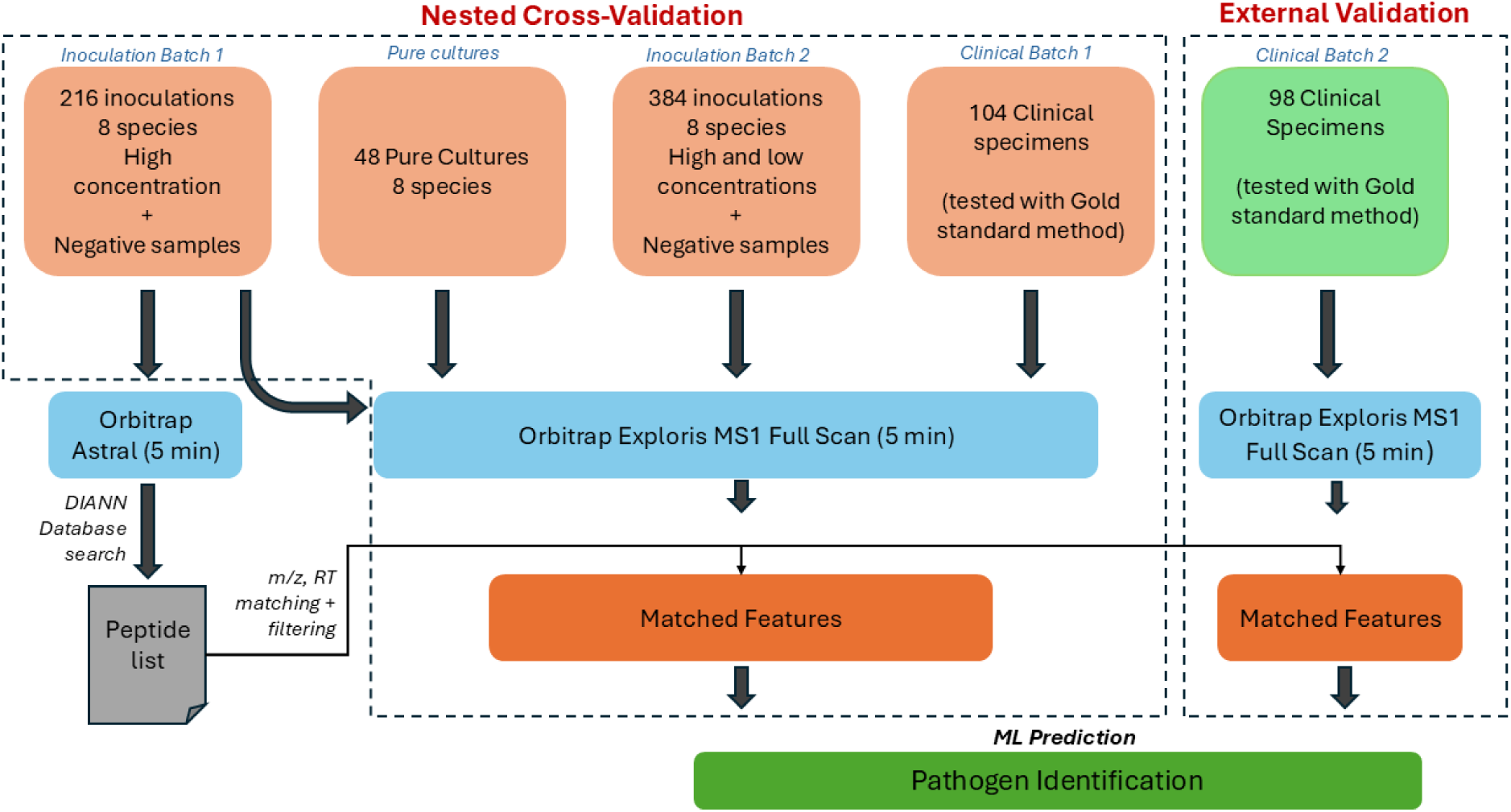
Illustration of overall workflow for peptide matching and machine learning. Three batches of data were used throughout the project in addition to pure bacterial cultures. The first two batches contained inoculated urine samples while the final batch contained clinical specimens. The first batch was analysed in DIA 5 min runs and the resulting DIA-NN database search results were used to establish the peptide list used for subsequent matching in MS1 only data. All subsequent batches were analysed only in MS1 Full scan mode and matched features were used as input for machine learning models, with inoculated batches, one clinical batch and pure cultures being used for training and testing while one batch of clinical samples were entirely withheld for final validation.

### Biomarker identification by DIA

The first batch of high-concentration inoculations were initially analysed via Data-independent acquisition (DIA) on the Orbitrap Astral mass spectrometer to determine a reference panel of candidate biomarkers for each of the eight target uropathogens. DIA analysis yielded a total of 8,218 candidate peptides following filtering to peptides consistently identified in bacterial samples with no identification in negative samples (**Supp.Table 3**). Each of the 8 bacterial species was represented by at least 200 unique peptides in the inoculated urine samples (**Supp.Fig 2**). The number of unique precursors per species was highest for *Pseudomonas aeruginosa* (2,901) and *Proteus mirabilis* (1,358), intermediate for *Escherichia coli* (953), *Klebsiella pneumoniae* (656), and *Staphylococcus saprophyticus* (611), and lower for *Staphylococcus aureus* (259), *Enterococcus faecalis* (225), and *Streptococcus agalactiae* (200). The remaining peptides were shared across multiple species. Patterns of peptide overlap reflected known phylogenetic relationships, with closely related species such as *E. coli* and *K. pneumoniae* sharing a high number of peptides, whereas taxonomically distant species such as *K. pneumoniae* and *S. aureus* shared none.

### Matched biomarkers in MS1-only data

To assess peptide detection in MS1-only mode, we applied our R-based precursor matching pipeline to all high-concentration samples. This pipeline matches MS1 signal to DIA peptides using m/z, retention time and isotopic cluster information for pairing. The first batch of inoculated urines, which had previously been analyzed by DIA to generate the candidate biomarker set, was reanalyzed using MS1-only acquisition. A second batch of inoculations and the clinical urine specimens were measured only in MS1-only mode. This approach allowed us to determine which DIA-derived candidate precursors could be consistently detected across all datasets and establish a robust feature set for machine learning. Application of the R-based precursor matching pipeline to samples resulted in 5,733 of the 8,218 DIA candidate precursors being detected in samples from the first batch analysed in Full Scan mode on Orbitrap Exploris 480. In samples from the second batch in Full Scan mode, 6,879 precursors were detected, while 5,769 were observed in patient samples from the first patient cohort and 3941 were identified in samples from the second patient cohort. A total union of 7,746 precursors were retained for downstream analyses (**Supp.Table 4**).

### Random Forest model training and feature utilization

Nine one-vs-all Random Forest classifiers were trained: one per target bacterial species and one for blank samples. All models achieved strong performance on the training data, with an average Matthews Correlation Coefficient (MCC) of 0.9938. Per-class MCC values were consistently high across all classes, each exceeding 0.98. *Pseudomonas aeruginosa* achieved a perfect MCC of 1.0 on the training data, followed closely by *Staphylococcus aureus* (0.996), blank samples (0.995), *Escherichia coli* (0.994), *Proteus mirabilis* (0.993), *Klebsiella pneumoniae* (0.992), *Streptococcus agalactiae* (0.991), *Staphylococcus saprophyticus* (0.990), and *Enterococcus faecalis* (0.985).

Across nested cross-validation iterations, the number of features retained per class-specific model was highly consistent. Mean feature set sizes ranged from 81.5 (Efa) to 95.4 (Ssa), with relatively low variability across runs (standard deviations ∼2.4–4.0 features). Minimum and maximum values were also stable, spanning 76–100 features depending on the class. Overall, most classes retained approximately 80–95 features per model, indicating stable feature selection under the consensus filtering procedure.

### Model performance across datasets

Model stability was evaluated using performance variability across the 20 outer cross-validation iterations **(Figure 3.a)**. Training MCCs were consistently high with a mean of 0.99 ranging from 0.988 to 0.998, while test MCCs ranged from 0.83 to 0.92 (mean 0.88), indicating robustness across splits. Per class performance was more variable ranging from 0.792 (Blank) to 0.967 (Pae), per class sensitivity was consistently above 0.80 ranging from 0.833 (Sau) to 0.963 (Pae), Per class specificity was very high consistently above 0.90 ranging from 0.949 (Blank) to 0.998 (Pae) and balanced accuracy was also consistently above 0.90 ranging from 0.910 (Efa) to 0.980 (Pae) **(Supp.Table 5).**

**Figure 3:**
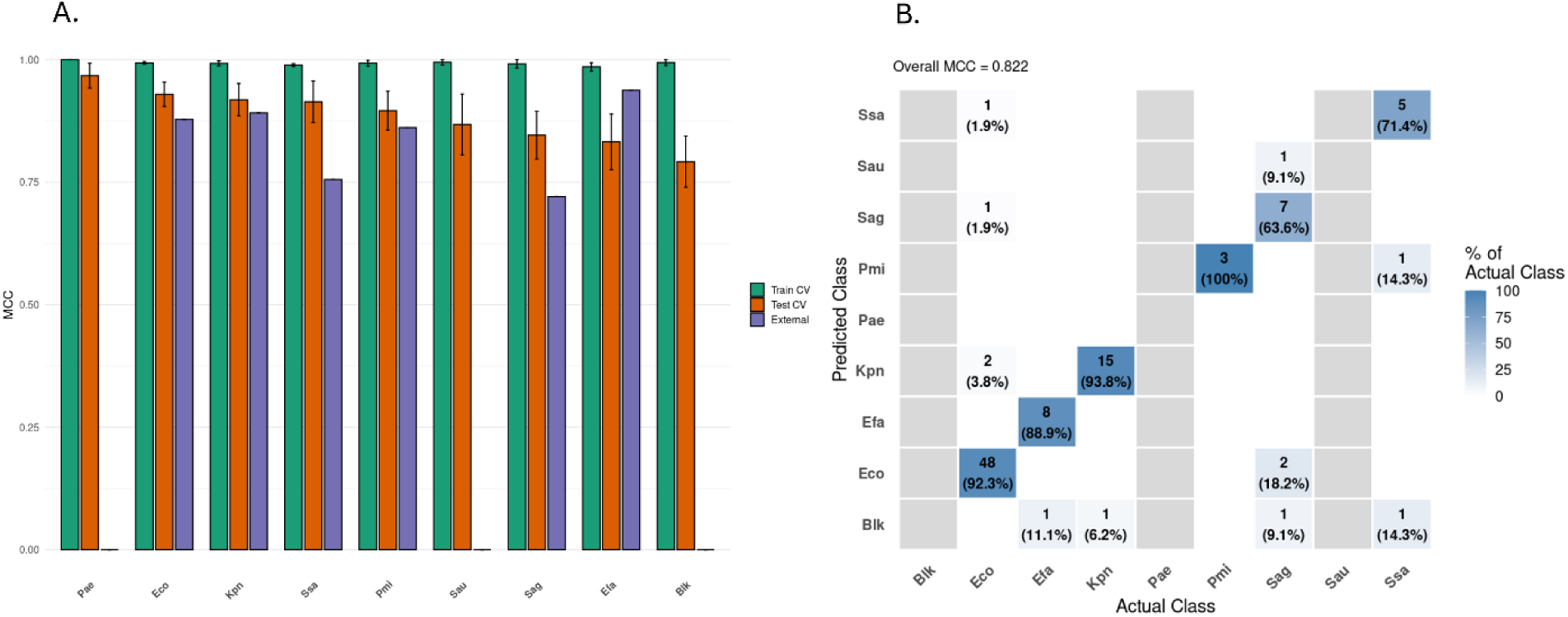
Detailed model performance on train and test splits in cross validation and on held-out patient dataset. **a. Matthews Correlation Coefficient (MCC) score evaluation on cross-validation outer train and test splits and evaluation of final trained model on external patient data** Bars in green represent the MCC score of each one vs all random forest model on training datasets (70% stratified split) and bars in red represent each model’s score on testing datasets (30% stratified split) and bars in purple represent performance on patient samples from a held-out batch for external testing. Class refers to each of the uropathogens and the positive class of each 1 vs All model: *Escherichia coli* (Eco), *Enterococcus faecalis* (Efa), *Klebsiella pneumoniae* (Kpn), *Pseudomonas aeruginosa* (Pae), *Streptococcus agalactiae* (Sag), *Staphylococcus aureus* (Sau), *Staphylococcus saprophyticus* (Ssa), *Proteus mirabilis* (Pmi), and blank (Blk, representing non-infected samples) **b. Detailed confusion matrix of predictions on patient dataset** The confusion matrix displays the agreement between the true class labels and the classes predicted by the model. Columns correspond to the actual (ground-truth) classes, while rows represent the classes predicted by the model. Each cell shows the number of samples assigned to a given predicted–actual class combination, along with the percentage relative to the total number of samples within the corresponding actual class. Percentages are visualized using a color gradient, where darker shading indicates a higher proportion of samples in that cell. Correct classifications occur along the diagonal of the matrix, where predicted and actual classes match. Off-diagonal cells represent misclassifications, indicating instances where samples belonging to one class were predicted as another. The Matthews correlation coefficient (Overall MCC) is reported as a summary metric of overall classification performance across all classes.

The model was then applied on a clinical batch containing 127 urine specimens never used for training or cross-validation and also analyzed with the gold standard method (culture + MALDI-TOF) (**Supp. Table 4**). When considering the 98 specimens found positive for one of the 8 target pathogens, our model achieved an overall MCC of 0.822, with correct classification of 86 samples (**Figure 3.b**). No *P. aeruginosa* or *S. aureus* infections were present in this sample set. For the six remaining species, per class MCCs consistently exceeded 0.70 ranging from 0.93 (*E. faecalis*) to 0.72 (*S. agalactiae*). Per class sensitivity was variable ranging from 0.64 (Sag) to 1 (Pmi), however sample counts are variable therefore metrics may be misleading (Pmi has the fewest samples at n=3), per class specificity was very high consistently above 0.95 ranging from 0.956 (Eco) to 1 (Efa) and balanced accuracy was also consistently above 0.80 ranging from 0.812 (Sag) to 0.995 (Pmi) **(Supp.Table 5).** Overall, pathogen-specific MCCs remained above 0.98 training, above 0.75 in testing and above (0.70) patient data, **(Figure 3.a)**.

### Predictions on edge cases

Samples inoculated with four species not in training (*Acinetobacter pittii* (Api), *Candida glabrata* (Cgl), *Serratia marcescens* (Sma), and *Providencia stuartii* (Pst)) at 10^6^ CFU/mL were misclassified, as expected for non-target pathogens of our classification method. Unknown species were often assigned to the most taxonomically similar known class: Sma mostly identified as *K. pneumoniae* (34/48), 37/48 Pst as *P. mirabilis*, Cgl predominantly as blank (22/48), and Api frequently as *P. aeruginosa (21/48)* or blank (19/48) (**Supp. Table 4**).

The general trend in classification of non-target pathogens in clinical samples seemed to also favor closest known species within the model e.g: *Klebsiella oxytoca* and *K. aerogenes* as *K. pneumoniae*. Coinfections with known pathogens only such as *E. Coli* and *P. mirabilis*, *P. mirabilis* and *E. faecalis*, *E. coli* and *E. faecalis* were predicted *E. Coli*, *P. mirabilis* and *E. Coli* respectively. However, with samples containing unidentified contaminants, results varied with some coinfections being correctly classified to the known class present, whilst others were assigned blank or other classes that may reflect the taxonomy of the unidentified contaminants (**Supp. Table 4**).

### Prediction confidence and score interpretation

Model prediction scores were compared across prediction to evaluate how the predicted class score can be used as an indicator of confidence **(Figure 4.a)**. A clear divide was observed between correct and incorrect predictions. Correct predictions consistently showed higher median scores: 0.858 in cross validation outer testing, and 0.960 in external patient samples. Incorrect predictions had lower medians: 0.224 (cross validation testing), 0.339 (external patient samples). samples containing non-target pathogens had median confidence of 0.253, reflecting increased ambiguity.

**Figure 4.**
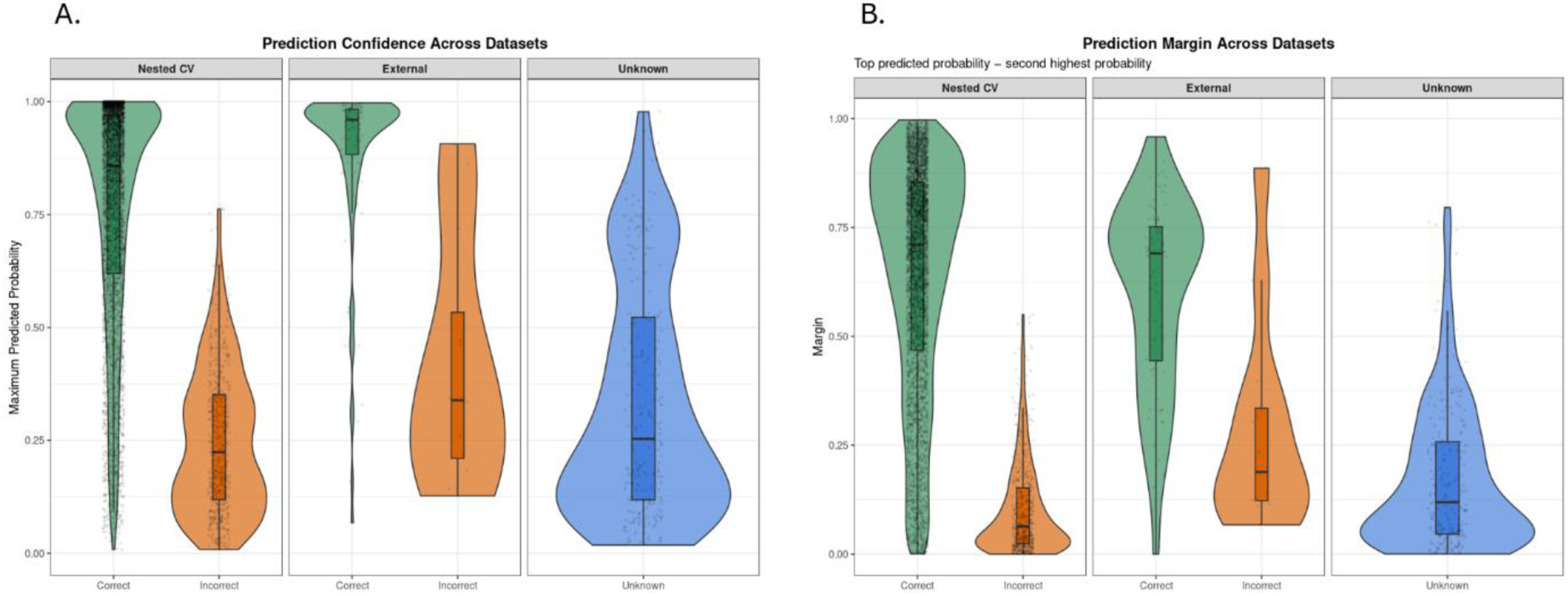
Model score comparison across correct and incorrect predictions on cross-validation testing, external patient testing and edge-case datasets. **a: Comparison of highest score of predicted class in correct and incorrect predictions across datasets.** Boxplots and violin plots represent the distribution of highest scores across predictions per dataset on cross validation test data, external patient validation and samples containing only non-target pathogens. Predictions are separated into correct classifications (green) and incorrect classifications (red) depending on whether the highest score corresponded to the actual class of the sample. Some datasets contain no correct predictions (Unknown) as they correspond to samples containing non-target pathogens that are not included in the model. **b. Comparison of margin between highest score and second highest score in correct and incorrect predictions across datasets.** Boxplots and violin plots represent the distribution of the margin between the top highest score and the second highest score across predictions per dataset on cross validation test data, external patient validation and samples containing only non-target pathogens. Predictions are separated into correct classifications (green) and incorrect classifications (red) depending on whether the highest score corresponded to the actual class of the sample. Some datasets contain no correct predictions (Unknown) as they correspond to samples containing non-target pathogens that are not included in the model.

Margin between top two predicted class scores further indicated confidence, with incorrect predictions associated with smaller margins **(Figure 4.b)**. A threshold of 0.2 effectively separated most correct from incorrect classifications; samples containing non-target pathogens had median margin of 0.119, reflecting increased ambiguity.

Although purely observational, these scores and thresholds help the user interpret the model’s output more effectively and better assess the confidence of a given prediction.

### Graphical user interface

An RShiny application was developed to integrate preprocessing and prediction. Users upload mzML files and a CSV of target peptides; a pre-configured CSV is provided. The app displays predictions with top-class, score, threshold indicator, and model-specific scores (**Supp.Fig 3**). Datasets ≤20 samples are processed sequentially; larger datasets are parallelized. Outputs can be downloaded as CSV, and the application is compatible with Windows and Linux.

## Discussion

In this study, we present a robust method for linking high-resolution data-independent acquisition (DIA) MS/MS data with high-resolution MS1-only data to identify and track bacterial biomarkers. This approach removes traditional limitations on the number of peptides analyzed and runtime, allowing rapid, high-throughput analysis of 300 samples per day, a throughput compatible with the requirements of clinical laboratories. The method was specifically applied to the classification of eight uropathogens found in 84% of urinary tract infections (UTIs) and demonstrated strong predictive performance, with Matthews Correlation Coefficient (MCC) scores of 0.88 on average on cross validation test data and 0.82 on patient samples correctly identifying the pathogen present in 86 out of 98 samples. These results highlight the potential of MS1-based workflows for clinical pathogen identification.

Despite this overall success, several limitations and areas for improvement were identified. Performance between training and testing splits during cross validation revealed a drop in MCC between training and testing with higher standard deviation on testing datasets. Per class MCCs showed higher variability with classes such as *P. aeruginosa* performing consistently well and other classes such as *S. agalactiae* and *E. faecalis* having more variability and lower MCCs. This may be linked to the lower number of individual peptide signals identified for these species which makes them more susceptible to variation across the cross validation splitting process where splits are stratified by class but may contain samples of varying concentrations. Negative samples (blank) were also classified with high variability and lower MCCs which is consistent with the method as we focus on features corresponding to signal detection in MS1 windows of expected bacterial signals therefore negative samples will have higher variation across samples. Despite this variability testing performance remained on average well above 0.80 across classes and still showed robust performance.

Finally, although our model performed well on patient data a small dip in performance was observed in per class MCC although *E. faecalis* achieved higher per class MCC than in cross validation training. This variation may be due to strain variability and urine heterogeneity as although 52 patient samples from a separate batch was included in training this batch was only a small part of the training data and was imbalanced and presented missing classes. Perfect balance across pathogen classes in such datasets is unlikely to be achievable as real-world clinical datasets are inherently imbalanced, with some pathogens (e.g., *E. coli*) far more prevalent than others. Inoculations in negative urine samples allow for more balanced training sets but may not reflect all the variability found in clinical samples. Despite these limitations overall MCC was 0.82 and per-class MCCs were consistently above 0.70 on the external clinical test dataset providing good overall classification.

Another challenge inherent to MS1-based classification lies in peptide identification and matching. Although our workflow relies on precise m/z values, retention time windows, and isotopic pattern filtering, some ambiguity persists and noise or interference may be detected as bacterial signals. Mitigation strategies include the use of a high number of biological replicates and broad biomarker panels in training, which help ensure that true species-specific signals are captured consistently. The MS1-only workflow is particularly well-suited for this strategy, as it allows rapid analysis of large sample sets while maintaining high resolution and sensitivity. Exploratory predictions on samples containing unknown pathogens reveal important considerations for clinical use. When predicting previously unseen pathogens or coinfections, the model tended to assign samples to the most taxonomically related species, reflecting shared peptide signatures however outliers persisted. In future work, expanding to additional pathogens may help mitigate these limitations and improve robustness under clinically relevant conditions.

In addition, by examining both prediction scores and margins between top predictions across cross validation, external clinical dataset performance and edge-case datasets, we show that small scores and low margins correspond to low model confidence, while high scores and large margins indicate reliable predictions. Considering the full probabilistic output in this way allows improved model interpretability, allowing users to gauge the reliability of individual predictions and easily flag potential errors. Exploiting these measures can support realistic evaluation and informed decision-making in clinical settings, using the machine learning predictions as a decision aid rather than a fully automated classifier.

Although the current workflow focuses solely on classification, preliminary results suggest that MS1 intensities could potentially support quantification, which provides important information to clinicians, as it can help guide the decision to initiate treatment depending on whether the specimen shows contamination (low concentration) or a true infection (high concentration). High-and low-concentration samples displayed consistent intensity differences for multiple biomarkers, indicating that quantitative relationships are preserved in MS1 data. Developing a robust quantification workflow would require additional experiments across a broader range of bacterial loads and careful calibration. Nonetheless, these initial observations are promising and suggest avenues for more detailed diagnostic applications in the future.

In conclusion, the combined DIA-to-MS1 matching approach provides a flexible, high-throughput method for biomarker identification and bacterial classification. While further optimization is needed to handle strain variability and unknown pathogens, this workflow establishes a foundation for rapid, species-specific diagnostics in urinary tract infections. Future developments could include expanded strain libraries, integration of quantitative MS1 data, and incorporation of prediction confidence metrics to enhance both accuracy and interpretability for clinical applications. Finally, although initially developed for UTIs, this approach has strong potential to be extended to other infectious diseases, additional clinical specimen types, and the detection of antimicrobial resistance.

## Data Availability

All code used for data processing and analysis necessary for prediction and use of the graphical interface is available on GitHub: https://github.com/Ellcoy/Urine_predict.

The raw mass spectrometry data are available on ProteomeXchange, on the following identifier: PXD075342.

## Ethics Statement

All urine specimens used were collected according to an ethical protocol approved by the Comité d’Éthique de la Recherche du CHU de Québec - Université Laval (recording number 2016-2656).

## Acknowledgments

We thank Milan Picard for helpful discussions and advice. We acknowledge Maciej Bromirski from Thermo Fisher Scientific for providing instrument support and technical expertise.

This work was supported by Genome Quebec (GQ139250), Thermo Fisher Scientific and Evosep Biosystems. Work was also supported by a scholarship from the Programme de bourses d’appui à la relève de l’Obvia – Observatoire international sur les impacts sociétaux de l’intelligence artificielle et du numérique. Graphical abstract created in BioRender. Roux-Dalvai, F. (2026) https://BioRender.com/8wuzara.

## Author Contribution

**Elloise Coyle :** Methodology, Software, Formal Analysis, Investigation, Validation, Visualization, Writing – Original Draft **; Antoine Lacombe-Rastoll :** Methodology, Investigation, Validation, Writing – Review and Editing**; Florence Roux-Dalvai :** Conceptualization, Supervision, Methodology, Validation, Writing – Review and Editing **; Mickaël Leclercq :** Conceptualization, Supervision, Methodology, Validation, Writing – Review and Editing **; Pascaline Bories :** Investigation, Writing – Review and Editing **; Clarisse Gotti :** Methodology, Investigation; **Ève Bérubé :** Resources **; Dorte Bekker-Jensen :** Resources **; Nicolai Bache :** Resources **; Sandra Isabel :** Resources **; Arnaud Droit :** Conceptualization, Supervision, Project administration, Funding acquisition

## Notes

https://github.com/Ellcoy/Urine_predict

### Competing Interest Statement

Dorte Bekker-Jensen and Nicolai Bache are employees of Evosep Biosystems

### Summary of Updates

Model training updated to include clinical samples, figures and supplementary data updated accordingly. Upgrade of model framework to nested cross validation.

