## Supplementary data for "Pairing Data Independent Acquisition and High-Resolution Full Scan for Fast Urinary Tract Infection Diagnosis"

**Supp.Table 1** Details of pathogens used for sample preparation

**Supp.Table 2** Uniprot protein databases for DIA analyses

**Supp.Table 3** Excel worksheet containing DIA peptide panel **See Excel file**

**Supp.Table 4** Excel worksheet containing input data for cross validation and model training and all model outputs on reported datasets **See Excel file**

**Supp.Table 5** Detailed metrics of model performance on Cross validation testing and on clinical specimens containing target pathogens

**Supp.Fig 1** Pipeline in R to process Full Scan mzML files and extract biomarker information

**Supp.Fig 2** Upset plot of Astral (DIA) biomarkers detailing unique and overlapping peptides between target bacterial species

**Supp.Fig 3** Screenshot of Graphical User interface of Rshiny app developed to make the predictive models and pre-processing steps readily available

**Supp.Table 1**. Details of pathogens used for sample preparation

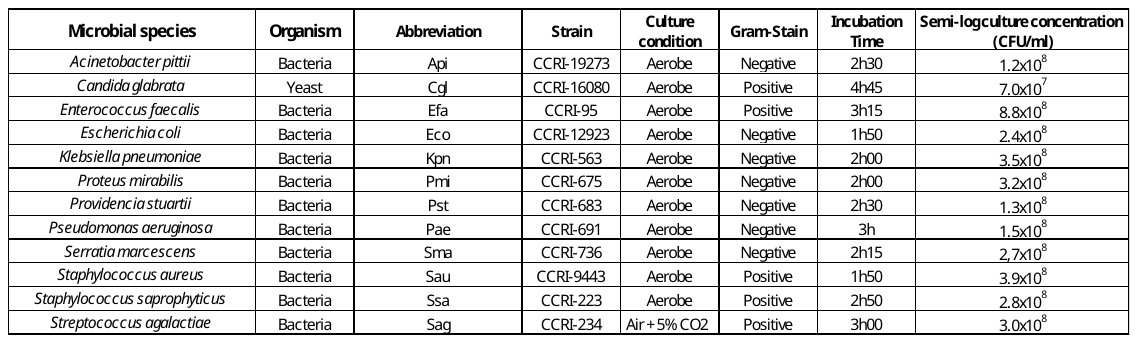

**Supp.Table 2**: Uniprot protein databases for DIA analyses

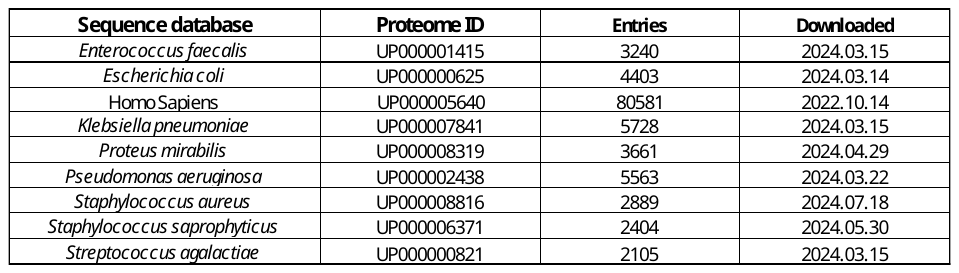

**Supp.Table 3** Excel worksheet containing DIA peptide panel **Please see Excel file**

**Supp.Table 4** Excel worksheet containing input training and testing data for feature selection and model training and all model outputs on reported datasets

**Please see Excel file**

**Supp.Table 5** Detailed metrics of model performance on Cross validation testing and on clinical specimens containing target pathogens

**Cross Validation Outer Test Performance**

| **Dataset** | **Class** | **MCC mean** | **MCC sd** | **Sensitivity mean** | **Sensitivity sd** | **Specificity mean** | **Specificity sd** | **Balanced Accuracy mean** | **Balanced Accuracy sd** |
| --- | --- | --- | --- | --- | --- | --- | --- | --- | --- |
| CV | Blk | 0.792 | 0.052 | 0.894 | 0.046 | 0.949 | 0.017 | 0.922 | 0.024 |
| CV | Eco | 0.929 | 0.025 | 0.927 | 0.040 | 0.992 | 0.006 | 0.960 | 0.019 |
| CV | Efa | 0.832 | 0.057 | 0.835 | 0.093 | 0.984 | 0.011 | 0.910 | 0.045 |
| CV | Kpn | 0.918 | 0.033 | 0.931 | 0.059 | 0.990 | 0.009 | 0.961 | 0.028 |
| CV | Pae | 0.967 | 0.025 | 0.963 | 0.032 | 0.998 | 0.003 | 0.980 | 0.017 |
| CV | Pmi | 0.896 | 0.040 | 0.888 | 0.062 | 0.992 | 0.006 | 0.940 | 0.030 |
| CV | Sag | 0.846 | 0.049 | 0.844 | 0.063 | 0.986 | 0.007 | 0.915 | 0.032 |
| CV | Sau | 0.868 | 0.062 | 0.833 | 0.076 | 0.993 | 0.006 | 0.913 | 0.039 |
| CV | Ssa | 0.914 | 0.042 | 0.909 | 0.058 | 0.993 | 0.006 | 0.951 | 0.029 |

**External Patient Dataset Performance**

| **Dataset** | **Class** | **MCC** | **Sensitivity** | **Specificity** | **Balanced_Accuracy** |
| --- | --- | --- | --- | --- | --- |
| *External* | *Blk* | *NA* | *NA* | *NA* | *NA* |
| External | Eco | 0.878131441 | 0.923076923 | 0.956521739 | 0.939799331 |
| External | Efa | 0.937556583 | 0.888888889 | 1 | 0.944444444 |
| External | Kpn | 0.891297997 | 0.9375 | 0.975609756 | 0.956554878 |
| *External* | *Pae* | *NA* | *NA* | *NA* | *NA* |
| External | Pmi | 0.861455317 | 1 | 0.989473684 | 0.994736842 |
| External | Sag | 0.72040873 | 0.636363636 | 0.988505747 | 0.812434692 |
| *External* | *Sau* | *NA* | *NA* | *NA* | *NA* |
| External | Ssa | 0.755507463 | 0.714285714 | 0.989010989 | 0.851648352 |

**Supp.Fig 1: Pipeline in R to process Full Scan mzML files and extract biomarker information**

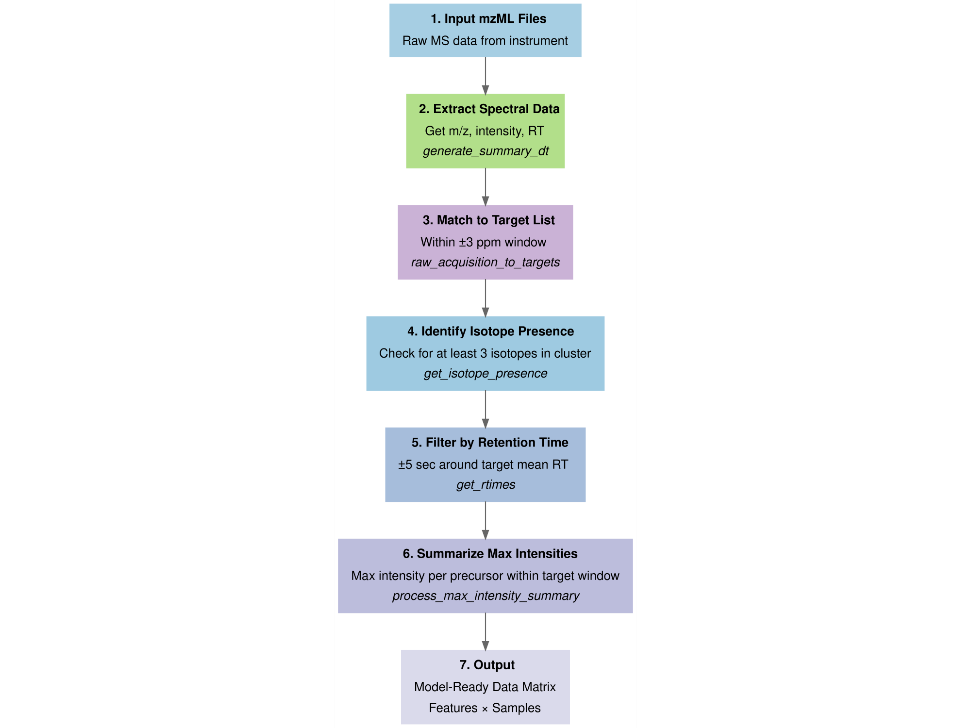
Each block corresponds to a step in the pipeline and names in italic refer to function names in the custom R code.

**Supp.Fig 2**. Upset plot of Astral (DIA) biomarkers detailing unique and overlapping peptides between target bacterial species

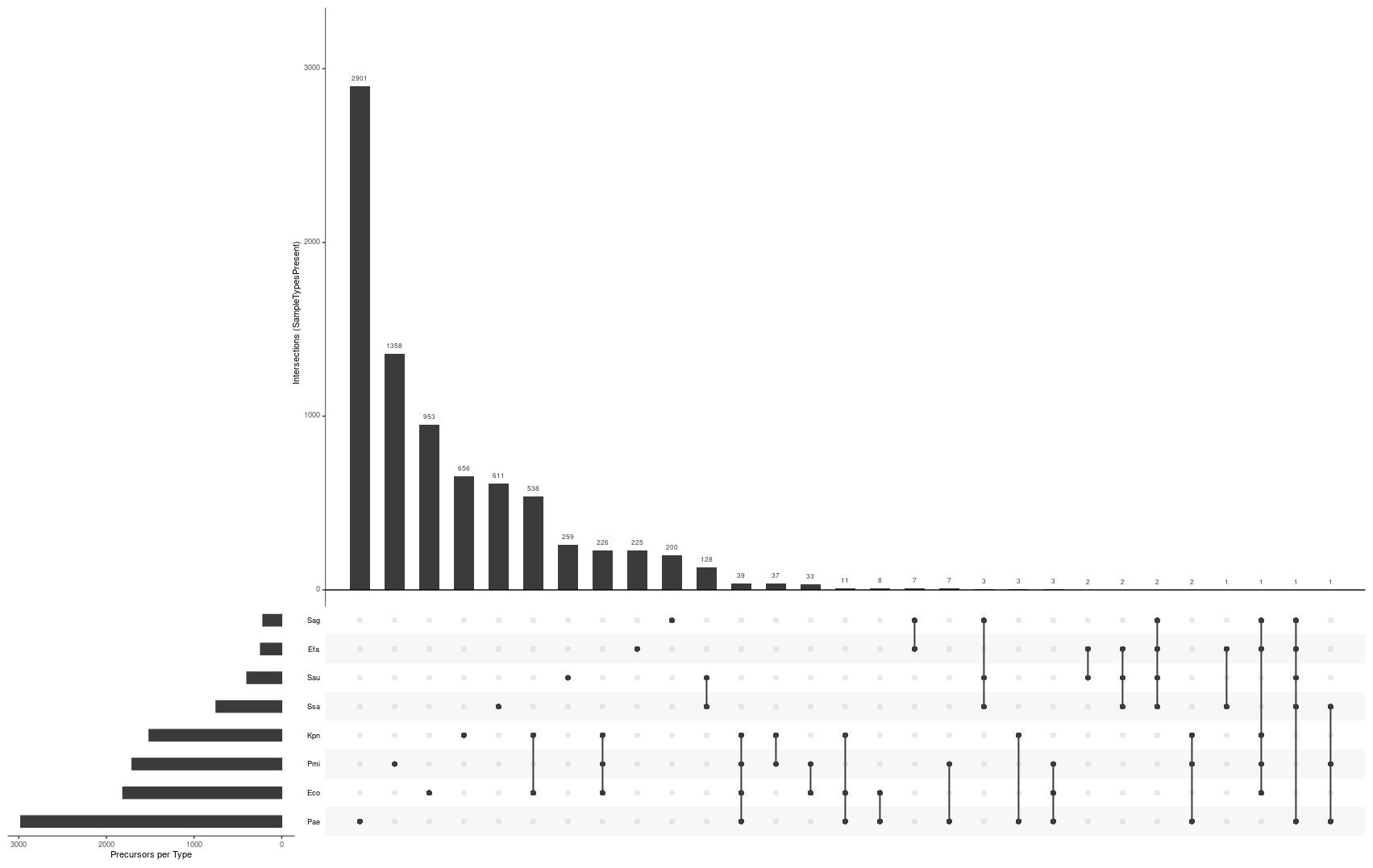

**Supp.Fig 3**. Screenshot of Graphical User interface of Rshiny app developed to make the predictive models and pre-processing steps readily available

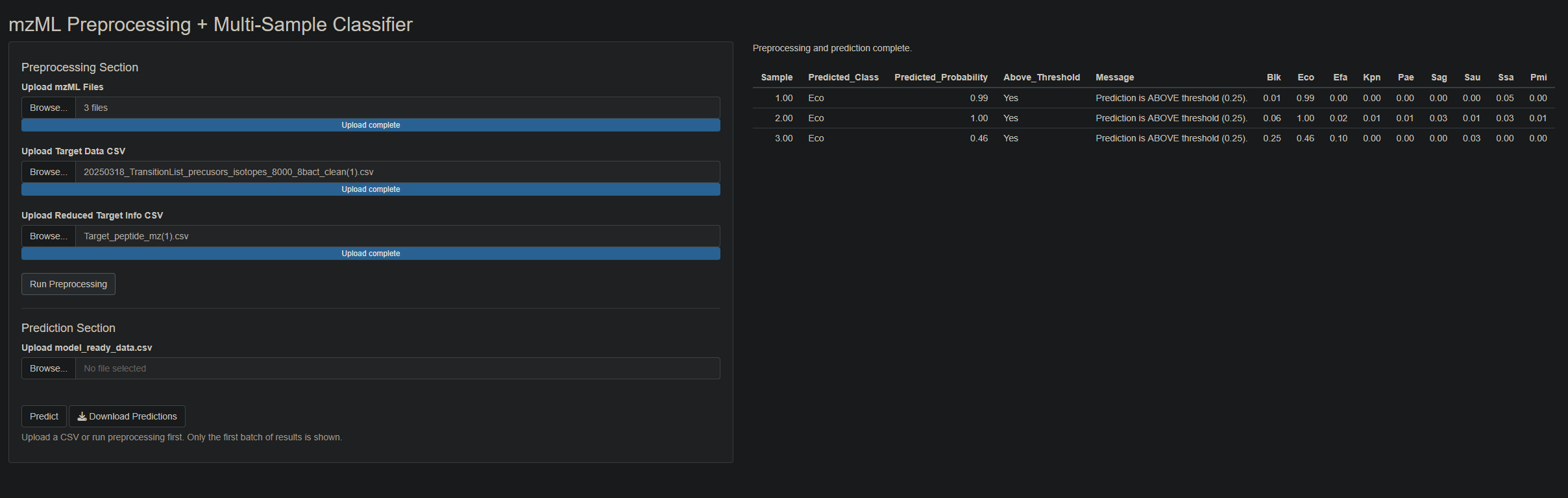
